# A reference panel of 64,976 haplotypes for genotype imputation

**DOI:** 10.1101/035170

**Authors:** Shane McCarthy, Sayantan Das, Warren Kretzschmar, Olivier Delaneau, Andrew R. Wood, Alexander Teumer, Hyun Min Kang, Christian Fuchsberger, Petr Danecek, Kevin Sharp, Yang Luo, Carlo Sidore, Alan Kwong, Nicholas Timpson, Seppo Koskinen, Scott Vrieze, Laura J. Scott, He Zhang, Anubha Mahajan, Jan Veldink, Ulrike Peters, Carlos Pato, Cornelia M. van Duijn, Christopher E. Gillies, Ilaria Gandin, Massimo Mezzavilla, Arthur Gilly, Massimiliano Cocca, Michela Traglia, Andrea Angius, Jeffrey Barrett, Dorret I. Boomsma, Kari Branham, Gerome Breen, Chad Brummet, Fabio Busonero, Hariy Campbell, Andrew Chan, Sai Chen, Emily Chew, Francis S. Collins, Laura Corbin, George Davey Smith, George Dedoussis, Marcus Dorr, Aliki-Eleni Farmaki, Luigi Ferrucci, Lukas Forer, Ross M. Fraser, Stacey Gabriel, Shawn Levy, Leif Groop, Tabitha Harrison, Andrew Hattersley, Oddgeir L. Holmen, Kristian Hveem, Matthias Kretzler, James Lee, Matt McGue, Thomas Meitinger, David Melzer, Josine Min, Karen L. Mohlke, John Vincent, Matthias Nauck, Deborah Nickerson, Aarno Palotie, Michele Pato, Nicola Pirastu, Melvin Mclnnis, Brent Richards, Cinzia Sala, Veikko Salomaa, David Schlessinger, Sebastian Schoenheer, P Eline Slagboom, Kerrin Small, Timothy Spector, Dwight Stambolian, Marcus Tuke, Jaakko Tuomilehto, Leonard Van den Berg, Wouter Van Rheenen, Uwe Volker, Cisca Wijmenga, Daniela Toniolo, Eleftheria Zeggini, Paolo Gasparini, Matthew G. Sampson, James F. Wilson, Timothy Frayling, Paul de Bakker, Morris A. Swertz, Steven McCarroll, Charles Kooperberg, Annelot Dekker, David Altshuler, Cristen Wilier, William Iacono, Samuli Ripatti, Nicole Soranzo, Klaudia Walter, Anand Swaroop, Francesco Cucca, Carl Anderson, Michael Boehnke, Mark I. McCarthy, Richard Durbin, Gonçalo Abecasis, Jonathan Marchini

## Abstract

We describe a reference panel of 64,976 human haplotypes at 39,235,157 SNPs constructed using whole genome sequence data from 20 studies of predominantly European ancestry. Using this resource leads to accurate genotype imputation at minor allele frequencies as low as 0.1%, a large increase in the number of SNPs tested in association studies and can help to discover and refine causal loci. We describe remote server resources that allow researchers to carry out imputation and phasing consistently and efficiently.

---

Over the last decade, large scale international collaborative efforts have created successively larger and more ethnically diverse genetic variation resources. For example, in 2007 the International HapMap Project produced a haplotype reference panel of 420 haplotypes at 3.1M SNPs in 3 continental populations^1^.More recently, the 1000 Genomes Project has produced a series of datasets built using low-coverage whole genome sequencing (WGS), culminating in 2015 in a reference panel (1000GP3) of 5,008 haplotypes at over 88M variants from 26 world-wide populations^2^. In addition, several other projects have collected low-coverage WGS data in large numbers of samples that could potentially also be used to build haplotype reference panels^3–5^. A major use of these resources has been to facilitate imputation of unobserved genotypes into genome-wide association study (GWAS) samples that have been assayed using relatively sparse genome-wide microarray chips. As the reference panels have increased in number of haplotypes, SNPs and populations, genotype imputation accuracy has increased, allowing researchers to impute and test SNPs for association at ever lower minor allele frequencies. A succession of methods developments have provided researchers with the tools to cope with these increasing larger panels^6–11^.

We formed the Haplotype Reference Consortium (HRC) (see **URLs**) to bring together as many WGS datasets as possible to build a much larger combined haplotype reference panel. By doing so, our aim is to provide a single centralized resource for human genetics researchers to carry out genotype imputation. Here we describe the first HRC reference panel that combines datasets from 20 different studies (**Supplementary Table 1**). The majority of these studies have low-coverage WGS data (4-8X coverage) and are known to consist of samples with predominantly European ancestry. However the 1000 Genomes Phase 3 cohort, which has diverse ancestry, is also included. This reference panel consists of 64,976 haplotypes at 39,235,157 SNPs that have evidence of having a minor allele count (MAC) greater or equal to 5.

We took the following approach to create the reference panel. We combined existing sets of genotype calls from each study to determine a ‘union’ set of 95,855,206 SNP sites with MAC >= 2. After initial tests, we decided for this first version of the HRC panel not to include small insertions and deletions (indels), since these were very inconsistently called across projects. We then used a standard tool to calculate the genotype likelihoods consistently for each sample at each site from the original study BAM files (see **Methods**) and make a baseline set of non-LD based genotype calls. We next applied a number of filters to remove poor quality sites (see **Methods**). We restricted this site list to sites with MAC >= 5 based on the calls made originally by the individual studies, corresponding to a minimum minor allele frequency (MAF) of 0.0077%, then added back sites that are present on several commonly used SNP microarray chips in GWAS. Sites with lower MAF would be likely to be poorly imputed. This site list consisting of 44,187,567 sites exhibited improved quality compared to the unfiltered MAC >=5 site list when assessed by measuring a per sample transition-to-transversion (Ts/Tv) ratio (**Supplementary Figure 1–2**). We also detected and removed 301 duplicate samples across the whole dataset (see **Methods**).

**Figure 1:**
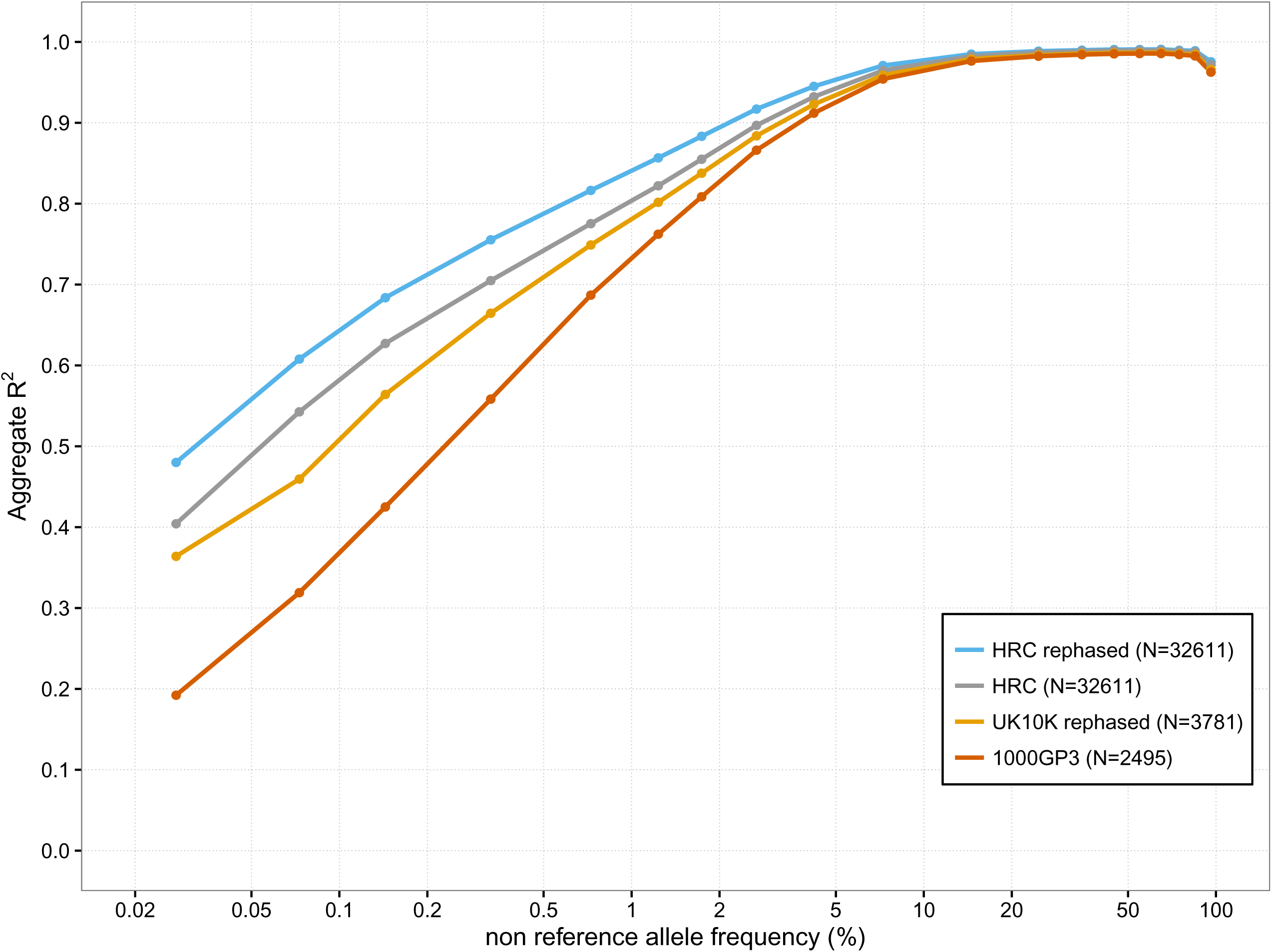
Performance of imputation using different reference panel. The x-axis shows the non-reference allele frequency of the SNP being imputed on a log scale. The y-axis shows imputation accuracy measured by aggregate r^2^ when imputing SNP genotypes into 10 CEU samples. These results are based on using genotypes from sites on Illumina OMNI 1M SNP array was used as pseudo-GWAS data.

**Figure 2:**
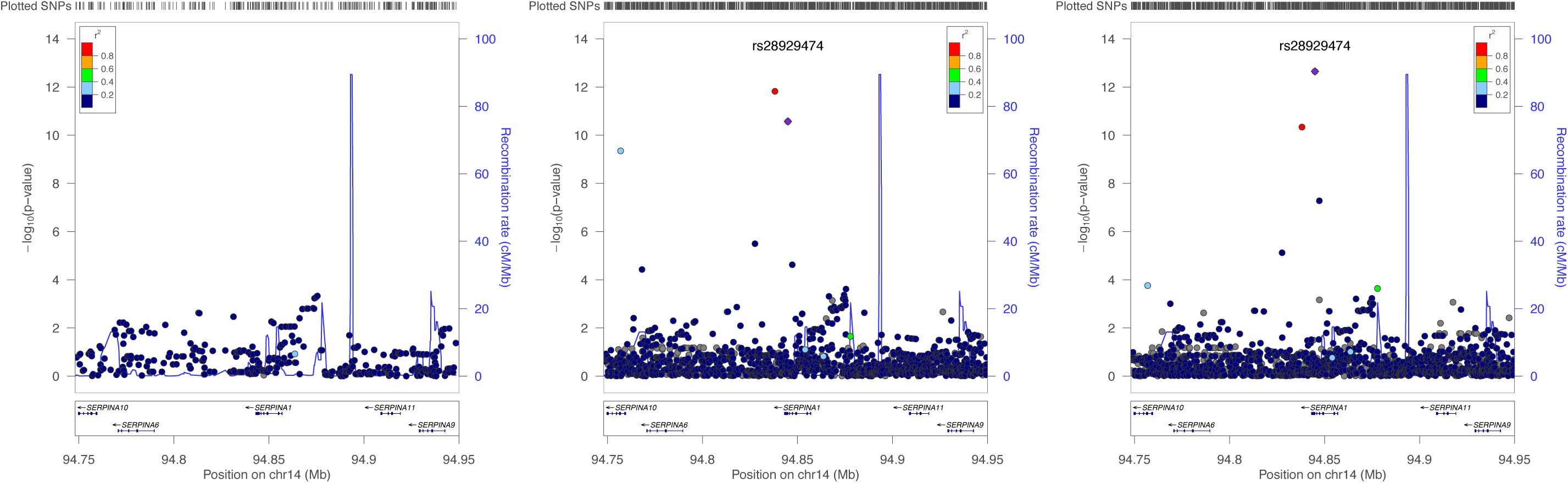
Association signal α_1_-antitripsin phenotype at the *SERPINA1* locus. Association test statistics on the-loglO p-value scale (y-axis) are plotted for each SNP position (x-axis). Three different imputation panels were used: HapMap2 (left), 1000GP3 (middle), HRC release 1 (right). The SNP rs28929474 is shown as a purple and other SNPs are coloured according to the levels of LD (r^2^) with this SNP (see r^2^ legend in each subplot).

Calling genotypes and phasing using low-coverage WGS data has been a computational challenging step for many of the 20 studies providing data. To reduce computation, we carried out this step on genotype likelihoods from all 32,611 samples together, and leveraged the original separately called haplotypes from each study to help reduce the search space of the calling algorithm (see **Methods**). We then applied a further refinement step by re-phasing the called genotypes using the SHAPEIT3 method^12^, based on experience from the UK10K project, which found this re-phasing approach produced substantially improved imputation accuracy when using the haplotypes^4^. After final genotype calling, we removed a further 123 samples (see **Methods**) and filtered out 4,952,410 sites whose MAC after refinement and sample removal was below 5, resulting in a final set of 39,235,157 sites and 32,488 samples. By measuring genotype discordance of the called genotypes compared to Illumina OMNI2.5M chip genotypes available on the 1000 Genomes samples we showed that both our site filtering strategy and the increased sample size of HRC led to improved accuracy (**Supplementary Table 2**). For example, we obtained a non-reference allele discordance of 0.39% on the full HRC dataset with site filtering, compared to 0.67% on the subset of 1000GP3 samples.

We next carried out experiments to assess and illustrate the downstream imputation performance compared to previous haplotype reference panels. To mimic a typical imputation analysis, we created a pseudo-GWAS dataset using high-coverage Complete Genomics (CG) WGS genotypes on 10 CEU samples (see **URLs**). We extracted the CG SNP genotypes at all the sites included on an Illumina 1M SNP array (Human1M-Duo v3C). These were used to impute the remaining genotypes which were then compared to the held out genotypes, stratifying results by MAF of the imputed sites. **Figure 1** shows that the HRC reference panel leads to a large increase in imputation performance when using a 1M SNP chip, compared to the 1000GP3 (R^2^=0.64 vs R^2^=0.36 at MAF = 0.1%) and also that the re-phasing step using SHAPEIT3 is worthwhile. HRC imputation at 0.1% frequency provides similar accuracy to 1000GP3 imputation at 0.6% frequency. **Supplementary Figure 3** and **4** show the results from a denser (Illumina OMNI 5M) SNP chip and a sparser (Illumina Core Exome).

To illustrate the benefits of using the HRC resource, we imputed a GWAS study of 1,210 samples from the InCHIANTI study^13^, including 534 that did not contribute to the HRC reference panel because they were not sequenced. Imputing using the HRC panel resulted in 15,501,516 SNPs passing an imputation quality threshold of r^2^≥0.5 compared to 13,238,968 variants (11,908,509 SNPs and 1,330,459 indels) when imputing using 1000 Genomes Phase 3, an increase of over 2 million variants. Taking the intersection of variant sites between the two panels to account for the filtering applied to the HRC panel resulted in 13,364,795 SNPs and 10,728,322 SNPs with r^2^≥0.5 for HRC and 1000 Genomes Phase 3 panel, respectively. The majority of these additional SNPs occur at the lower frequency range (**Supplementary Table 3**).

We next tested the HRC imputed genotypes for association with 93 circulating blood marker phenotypes, including many of relevance to human health such as lipids, vitamins, ions, inflammatory markers and adipokines^14,15^.This analysis highlighted potential novel associations at the nominal GWAS significance threshold of 5e-8 (**Supplementary Table 4**). When we repeated the imputation using the HRC panel without the overlapping InCHIANTI samples, we obtained similar results (**Supplementary Table 4**). We took these SNPs forward for replication in the SHIP and SHIP-TREND cohorts (see **Methods**) and found that two of the SNPs replicated (**Supplementary Table 5**). Specifically, we found that SNP rs150956780 (MAF= 0.6%) was associated with the Lactic Dehydrogenase phenotype (meta-analysis *p*-value = 3.779E-29) and SNP rs147142246 (MAF= 0.6%) was associated with the Potassium phenotype (meta-analysis *p*-value = 8.7E-09). We also found thatitis possible for HRC imputation to refine signals of association. For example, **Figure 2**shows the association results of HapMap2,1000GP3 and HRC based imputation for the a iantitripsin phenotype at the *SERPINA1* locus. HRC imputation gives a clear refinement of the signal at the rare causal SNP rs28929474 (MAF=0.5%) (**Supplementary Table 6**), known to predispose to the alpha 1 antitrypsin deficiency lung condition emphysema^16,17^. Similar results were obtained when using the HRC panel that excluded the InCHIANTI samples (data not shown).

Since the HRC reference panel combines data from many different studies with a range of restrictions on data release we have developed centralized imputation server resources (see **URLs**). Under this model researchers upload phased or unphased genotype data and imputation is carried out on central servers. Once completed researchers can download imputed datasets. Along similar lines, we have also developed a lower throughput phasing server for haplotype estimation of clinical samples with genotypes from high-coverage WGS data that takes advantage of rare variant sharing^18^ (see **URLs**). A limited subset of HRC haplotypes will be made available for researchers via the European Genomephenome Archive (EGA) for the sole purpose of phasing and imputation.

This first release of the HRC is the largest human genetic variation resource to date and has been created via an unprecedented collaboration of data sharing across many groups. We envisage continuing to expand the HRC and are currently planning a second HRC release differing from the first release in two ways. Firstly, we aim to substantially increase the ethnic diversity of the panel, by including data from sequencing studies in world-wide sample sets such as the CONVERGE study^19^, AGVP^20^ and HGDP^21^ Secondly, we aim to include short insertions and deletions in addition to SNP variants. In the limit of a reference panel consisting of the whole human population except the person being imputed, then imputation would likely be almost perfect for alleles at any frequency, since the panel would contain close relatives that share long and almost identical tracts of sequence. Therefore, we do expect to be able to make future gains in imputation performance. In some populations that have experienced isolation (like Sardinia or Iceland) we expect to approach this limit much faster. Thinking further ahead, we hope to work closely with efforts under way to collect large samples of high-coverage sequenced samples such as the UK 100,000 Genomes Project (see **URLs**).

## Online methods

### Union site list

Every study provided us with their most recent version of their haplotypes in VCF format with one VCF for every autosome. For every cohort, bcftools (v0.2.0‐rc12) was used to create an entire-autosome, SNP-only site list with alternate and total allele count information from these per-chromosome haplotypes. Multiallelic SNPs were broken into biallelics using ‘bcftools norm’. These per-cohort site lists were merged into a single file using an in-house Perl script that correctly merges alternate and total allele counts. We created site lists called MAC2 and MAC5 containing only sites with a minor allele count (MAC) across all studies of >= 2 and >=5, respectively, using bcftools. These sites lists contained 95,855,206 and 51,060,347 sites, respectively.

### Genotype likelihood calculations

The 'samtools mpileup' command was used to generate genotype likelihoods (GLs) at all MAC2 sites on a per sample basis from each sample’s BAM file. The pipeline and software versions have been made available online (see **URLs**). The resulting BCF files were merged using the 'bcftools merge' command and the MAC2 sites and alleles extracted using the 'bcftools call' command. The use of 'bcftools call' here made a baseline set of non-LD based genotype calls for each site across all samples. These calls were used for some initial sample QC (see Sample filtering section). We calculated GLs on 33,070 samples in total.

### Site filtering

We used an ad-hoc method for initial variant filtering which enabled us to identify variants that had been filtered out ‘quite often’ by our submitting studies. For each site and for each cohort, we labelled the site as “called” in that study if the putative calls from bcftools based on GLs exhibited more than one allele in that cohort, or “not called” if it showed no variation. We also used the haplotype sets provided by each study to determine whether each study had filtered out each site or not using their own internal calling pipeline. To determine a threshold of “number of times filtered out”, we stratified the sites according to their called status versus their filtered status (**Supplementary Figure 5**). We also measured the Ts/Tv ratio of the set of SNPs for each of these stratified combinations. SNPs corresponding to the cells above the red line in the figure were filtered out, removing all cells which had been filtered out by more than 4 studies or have Ts/Tv ratio less than 1.7.

We also applied a set of additional site filters as follows. We filtered out sites not on the MAC5 site list to restrict the site list to those that could be imputed well. We also filtered out sites if (i) any study (apart from 1000 Genomes) had a Hardy-Weinberg Equilibrium (HWE) *p*-value < 10^−10^, (ii) any study (apart from 1000 Genomes) had an overall inbreeding coefficient < −0.1, (iii) a MAF>0.1 with the site being called in fewer than 3 of the studies and not called in 1000 Genomes (the latter restriction kept sites present at high frequencies in non-European populations that were only called in 1000 Genomes). We also filtered out sites called only in the GoNLstudy or IBD cohort. We completely excluded GPC haplotypes from this step of the site list creation process.

After applying these filters, the site list comprised of 44,038,997 sites. Finally, we made sure that 4,914,335 sites found on a selection of common SNP genotyping arrays and those used in the GIANT consortium and the Global Lipids Consortium (**Supplementary Table 7**) were included in the final site list. The final site list after this filtering contained 44,187,567 sites.

### Sample filtering

Having used 'bcftools call' to extract sites and alleles, we had a set of baseline non-LD genotype calls (see Genotype likelihood calculations section). Based on these calls for chromosome 22, some outlier samples were evident and we removed 150 samples showing evidence for fewer than 10,000 non-reference SNPs or more than 10 singletons across the chromosome. This left a total of 32,920 samples.

To detect possible duplicates we used the original genotype calls submitted by the individual studies. We selected 1000 random sites that (1) were biallelic; (2) had European minor allele frequency > 5% in 1000GP3; and (3) had no missing data in any of the individual studies. Using the 'bcftools gtcheck' command, we counted the number of genotypes that differed between each sample pair. There was a clear set of 269 sample pairs with very few genotypes differing over the 1000 sites. We identified these samples as duplicates either within or between studies and removed one of the samples in the pair as described in **Supplementary Table 8.** Due to some samples being represented more than twice, there were a total of 261 samples removed due to duplicates. Before genotype calling, we also removed (i) 9 samples for which we had Complete Genomics data so that we could use these samples for testing purposes, (ii) 31 samples from 1000GP3 that were related samples (see **URLs**), (iii) 8 samples from the HELIC, AMD and ProjectMinE studies with sample labeling inconsistencies. These filters resulted in 32,611 samples being used for the genotype calling and phasing steps.

In addition, after the phasing, 83 samples from the AMD study were removed as the consent for these samples had been removed. We also repeated the duplicate detection process on the final HRC genotype calls, since some studies increased in size late on within the analysis process. This resulted in an extra 40 samples being removed and a total of 32,488 samples in the final phased reference panel.

### Genotype calling method leveraging existing haplotype calls

We called genotypes from the genotype likelihoods computed on the HRC samples by extending the SNPTools^22^ algorithm to leverage pre-existing haplotypes available from each cohort. Like other phasing and calling approaches^8,10^, SNPTools is an MCMC approach in which each sample's haplotypes and genotypes are iteratively updated using the current estimates of all other samples. A low-complexity Hidden Markov Model (HMM) with just four states is used to update each sample, where the states are a set of four "surrogate parent" haplotypes. The MCMC sampler employs a Metropolis-Hastings (MH) step to sample the set of surrogate parents. In large sample sizes the search space for these surrogate haplotypes is huge and results in low acceptance rates for the sampler. Our extension, called GLPhase (see **URLs**) uses pre-existing haplotypes to restrict the set of possible haplotypes from which the MH sampler may choose surrogate parent haplotypes. For each individual, we restrict the search space to 200 haplotypes that most closely match the two pre-existing haplotypes of the individual using a Hamming distance metric (100 for each haplotype). We run the method on chunks of 1,024 sites at a time, which is the default setting for SNPtools. Since the pre-existing haplotypes from each study do not contain exactly the same set of sites we filled in missing alleles in the preexisting haplotypes at our site list using the major allele at each site.

Restricting the search space in this way allows us to reduce the number of burnin iterations from 56 to 5, the number of sampling iterations from 200 to 95, and the number of MH steps taken at each iteration for each individual from 2*N* to 100, where *N* is the number of samples being phased. This reduces the complexity of our phasing algorithm from *O*(*N*^*2*^) to *O*(*N*). Although our implementation of the Hamming distance search has complexity *O*(*N*^*2*^), for *N* = 30,000, the impact of the search on run time is small (~5% of run time on each chunk). A chunk of 1024 sites can be phased in ~200 minutes using ~1.3GB of RAM. Once sample sizes are encountered where the Hamming distance search begins to dominate, our implementation could be replaced with *O*(*N log N*)clustering algorithms that we have implemented within the SHAPEIT3 algorithm^12^.

To illustrate how important GLPhase was to genotype calling and phasing on such a large sample size, we carried out a comparison to Beagle 3.1, Beagle 4.1 and the original SNPTools method. We ran all four methods on five randomly selected 1024 site chunks from chromosome 20 on the cluster using increasing sample sizes and measured run time. **Supplementary Figure 6** shows that GLPhase is approximately 100 times faster than the next quickest method at the full HRC sample size.

### Final phasing and haplotype estimation

We estimated haplotypes from GLPhase genotype calls using SHAPEIT3^12^. Chromosomes were phased in chunks consisting of 16,000 variants plus 3,300 variants overlapping with neighboring chunks on either side. The non-default command line option-w 0.5 was used for SHAPEIT3. Chunks were ligated using the ligateHAPLOTYPES program (see **URLs**). SHAPEIT3 does not handle multiple variants at the same genomic coordinate, so multiallelic sites (SNPs with 3 or 4 alleles) were shifted by one or two base pairs for rephasing, and then moved back to their original position after chunk ligation.

### Evaluation of genotype calling process

We tested the genotype calling process on data from chromosome 20 with different combinations of site lists and sample sets to assess both the effects of site filtering and the benefits of increasing samples size. We evaluated 3 different site lists: the 1000 Genomes Phase 3 set of sites (775,927), our HRC MAC5 site list (1,128,114) and our HRC MAC5 site list with additional site filtering (1,006,559). We ran the genotype calling method on 3 different sets of samples: the 2,525 original 1000 Genomes Phase 3 samples, a subset of 13,309 HRC samples that we used at an early stage of HRC testing (HRC Pilot) from studies 1000GP3, AMD, GoNL, GoT2D, ORCADES, SardinIA, FINLAND and UK10K, and the near-final full set of 32,905 HRC samples. We called genotypes using GLPhase on each of these 9 datasets and examined genotype discordance compared to Illumina OMNI2.5M genotypes produced by the 1000 Genomes Project. For this comparison, we focused only on genotypes from 365 samples shared across the 3 sample sets and at 42,244 SNP sites. We calculated percentage discordance for the 3 possible genotypes consisting of reference (REF) and alternate (ALT) alleles as well as an overall non-reference allele discordance rate (NRD). Results are shown in **Supplementary Table 2.**

### Downstream imputation performance

We assessed imputation accuracy of 4 different reference panels: 1000 Genomes Phase 3, UK10K, and two versions of the HRC reference panel, with and without re-phasing with SHAPEIT3. To do this we used high-coverage WGS data made publicly available by Complete Genomics (CG) (see **URLs**). For the pseudo-GWAS samples we used data from 10 CEU samples that also occur in the 1000 Genomes Phase 3 samples. These samples were removed from the various reference panels before using them to assess imputation performance.

Three pseudo-GWAS panels were created based on three chip lists (see **URLs**):The Illumina Omni 5M SNP array (HumanOmni5-4vl-l_A), the Illumina Omni 1M SNP array (Human1M-Duo v3C), and the Illumina Core Exome SNP array (humancoreexome-12vl-l_a). For these comparisons we only used sites in the intersection of the reference panels to enable a direct comparison.

These pseudo-chip genotypes were used to impute the remaining genotypes which were then compared to the held out genotypes, stratifying results by MAF of the imputed sites.

Imputation was carried out using IMPUTE2^7^ which chooses a custom reference panel for each study individual in each 2 Mb segment of the genome. We set the khap parameter of IMPUTE2 to 1000. All other parameters were set to default values. We stratified imputed variants into allele frequency bins and calculated the squared correlation between the imputed allele dosages at variants in each bin with the masked CG genotypes (called aggregate r^2^ in **Figure 1**). Nonreference allele frequency for each SNP was calculated from HRC release 1 GLs at MAC>=5 sites. **Figure 1** shows the results for the Illumina Omni 1M chip. **Supplementary Figures 3** and **4** show the results from the Illumina Core Exome chip and the Illumina Omni 5M chip respectively.

### Details of imputation, association testing and replication in the InCHIANTI study

A total of 1,210 individuals from the InCHIANTI study were genotyped using the Illumina Infinium HumanHap550 genotyping array^13,14^. Individuals were prephased using autosomal SNPs after filtering out SNPs with MAF <1%, Hardy-Weinberg *p*-value <10 ^−04^, and missingness >1%. SNPs were also removed if they could not be remapped to the GRCh37 (hg19) human reference. This resulted in 483,991 SNPs available for pre-phasing. Phasing was performed locally using SHAPEIT2 ^10^.

Imputation was performed remotely using the Michigan Imputation Server (see **URLs**). A total of 39,235,157 SNPs and 47,045,346 variants were imputed from the HRC and 1000 Genomes Phase 3 (v5) reference panels, respectively. An imputation quality threshold of r^2^ >0.5 was subsequently applied to both imputation datasets prior to association testing. This resulted in 15,501,516 and 13,589,949 variants available for association analysis derived from HRC‐ and 1000 Genomes-based imputation, respectively.

A total of 93 circulating factors available in the InCHIANTI study were double inverse-normalised, while adjusted for age and sex, prior to association testing ^14^’^15^. Association analysis was performed using a linear mixed model framework as implemented in GEMMA (see **URLs**). Plots of association in **Figure 2** were produced using LocusZoom (see **URLs**).

We attempted to replicate the associations reported in Supplementary Table 3 in the SHIP and SHIP-TREND cohorts^23^. The SHIP samples were genotyped using the Affymetrix Genome-Wide Human SNP Array 6.0. The SHIP-TREND samples was genotyped using the Illumina Human Omni 2.5 array. Prior to imputation, duplicate samples (by IBS), samples with reported vs. genotyped gender mismatch or samples with a very high heterozygosity rate were excluded. Additionally, all monomorphic SNPs, SNPs with duplicate chromosomal position, SNPs with pHWE <0.0001 and SNPs with a callrate <95% were filtered. Imputation was performed on the Sanger Imputation Service (see URLs) against the HRC panel. In total, 4,070 SHIP samples and 986 SHIP-TREND samples were included in the imputation of genotypes. Association analyses were conducted using SNPTEST v2.5.2^24^.

## URLs

Haplotype Reference Consortium

http://www.haplotype-reference-consortium.org/

Michigan Imputation Server

https://imputationserver.sph.umich.edu/

Sanger Imputation Server

https://imputation.sanger.ac.uk/

Oxford Phasing Server

https://phasingserver.stats.ox.ac.uk/

Genotype Likelihood calculation scripts

https://github.com/mcshane/hrc-releasel

GLPhase

http://www.stats.ox.ac.uk/~marchini/software/gwas/gwas.html

ligateHAPLOTYPES

https://mathgen.stats.ox.ac.uk/genetics_software/shapeit/shapeit.html

Complete Genomics high-coverage WGS genotypes

http://ftp.1000genomes.ebi.ac.uk/voll/ftp/technical/working/20130524_cei_combined_calls/

1000 Genomes Project OMNI genotypes

ftp://ftp.1000genomes.ebi.ac.uk/vol1/ftp/release/20130502/supporting/hd_genotvpe_chip/ALL.chip.omni_broad_sanger_combined.20140818.snps.genotypes.vcf.gz

100,000 Genomes Project

http://www.genomicsengland.co.uk/the-100000-genomes-proiect/

GEMMA

http://www.xzlab.org/software.html

LocusZoom

http://locuszoom.sph.umich.edu/locuszoom/

1000GP3 related samples

ftp://ftp.1000genomes.ebi.ac.Uk//voll/ftp/release/20130502/20140625_related_individuals.txt

SNP chip site lists

http://www.well.ox.ac.uk/~wrayner/strand/

## Acknowledgements

J.M acknowledges support from the ERC (Grant no. 617306). W.K acknowledges support from the Wellcome Trust (Grant no. WT097307). S.M and R.D acknowledge support from Wellcome Trust grant WT090851. We are grateful to all participants of all the studies that have contributed data to the HRC.

## Author contributions

The HRC was initially conceived by discussions between J.M, G.A, R.D, M.M and M.B. Analysis and methods development was carried out by S.M, S.D, W.K, O.D, A.R.W, P.D, H.K. Supervision of the research was provided by J.M, G.A and R.D. The Michigan Imputation server was developed by C.F, L.F, S.S and G.A. The Sanger Imputation server was developed by P.D, S.M and R.D. The Oxford Statistics Phasing server was developed by W.K, K.S and J.M. All other authors contributed datasets to the project or provided advice.

